# Clinico-Demographic trend of HIV-positive cases and sero-discordance at a secondary level hospital in Haryana, North India-programmatic implications for a low HIV prevalence State

**DOI:** 10.1101/356881

## Abstract

**Background:** Appropriate programmatic intervention for HIV Care and Treatment in a low prevalence state requires local level analysis of programme data. Data generated at an Integrated Counseling and Testing Centre (ICTC) may provide crucial information to understand the epidemiology of the disease in a particular region. There is paucity of information on HIV epidemiology at sub-district level in a low HIV prevalence State of India.

**Methods:** A secondary analysis of the records from January to December for the years, 2009 through 2014 was conducted among clients who tested HIV positive at the ICTC of a sub-district hospital in Haryana, North India.

**Results:** A total of 199 individuals were tested HIV positive of whom 121 (61%) were males. By age-group, 8, 8, 178, and 5 individuals were respectively in <5, 5-18, 18-59 and >60 years of age. Over years from 2009 through 2014, 11, 12, 30, 37, 51 and 58 people tested HIV positive, with no sigfinicant sex difference (chi2 p =0.929). Statistically non-significant increase of 18-59 years individuals was observed, from zero in 2009 to 5 in 2014. Major route of transmission was heterosexual (80%), followed-by, parent-to-child (5%), Blood Transfusion (1.5%), MSM (1%) and FSW (0.5%). One third each, were self-referred, from government facility; 16% from tuberculosis clinic. Median CD4 count in 2014, was 392. Serodiscordance rate spouses of HIV positive females was 17%, of males was 33%.

**Conclusion:** Analysis of programme data at a sub-district ICTC could highlight emerging trend even in a low HIV prevalence state.

## Introduction

Significant progress has beeen made in halting and reversing the HIV epidemic in India. By 2015, The evidence based strategic approach for prevention and control of HIV epidemic in India has resulted in a 66% decline of new HIV infections from 2000.(1) Despite these dramtic and exemplary efforts led by India’s National AIDS Control Organization, an estimated 85,000 (56,000 to 120,000) new HIV infections would have occurred in 2015, of which, almost 90% are among those more than 15 years of age. (1)

With the exception of mother to child transmission, and those transmission scenarios related to blood products, all new infection must be due to some sort of discordance between partners ie where one is HIV positive and the other HIV negative. HIV sero discrodance could be of following types-among monogamous married couples when either male or female is HIV positive; among men who have sex with men in which case, the index case could transmit to potentially more than one partner depending upon whether the person is bisexual, male sex worker, or uses injecting drugs; among male injecting drug users transmitting to fellow IDUs, and sexual partners; among females IDUs, transmitting to those sharing needles with them, and their sexual partners.(2)

HIV negative individuals in discordant partnerships are at high risk of infections and preventing infections targeted at such individuals is crucial.(3) The intervention could be targeted to the uninfected partner, thereby minimizing risk of acquisition (Pre Exposure Prophylaxis); could be targeted too infected partner so as to control risk of transmission e.g through anti retro viral therapy (ART); or to both the partners through a common intervention such as use of condom, or use of safe needles and syringes. (3)

Limited availble information from studies done at tertiary care, teaching hospitals indicate a sero-discordance rate of more than 40%. Tertiary care teaching hospitals are not representative of the population of the district due to the care seeking pattern. However, no information is available on the quantum of HIV sero-discordance among married couples avaialing HIV testing services at sub-district leve hospitals. in India. We did this study to determine the level of sero-discordance among HIV positive partners at an integrated counseling and testing centre of a sub district level hospital in Faridabad district, Haryana, north India.

## Methods

*Study design and setting–* The study was based on analysis of routine administrative health data collected at the Integrated Counseling and testing Centre (ICTC) for HIV, situated at a sub district hospital of Ballabgarh, in the Northern Indian State of Haryana. This ICTC is a type of facility intergrated HIV counseling and testing centre. The ICTC follows the guidelines issued by the National AIDS Control Organization, India.(4) The ICTC catered to direct walk-in clients, women visiting the hospital for antenatal checkup, and suspected tuberculosis patients referred from the Tuberculosis Unit at the same hospital.

The sub district hospital is situated in Ballabgarh town, on the outskirts of district of Faridabad in the Northern State of Haryana, in India. The nearest district hospital is at a distance of around 8 kms, the district Faridabad. The hospital provided daily outpatient services in the disciplines of Medicine, Surgery, Obstetrics and Gynecology, Pediatrics, Ophthalmology, Dentistry and Ayurveda (Indian System of Medicine). This hospital is under the administration of the Centre for Community Medicine, All India Institute of Medical Sciences, New Delhi, in collaboration with the Haryana Governement, and forms an integral care component for the Ballabgarh HDSS.(5)

*Data Management and Analysis–* The data was abstracted for those clients who tested HIV positive at the ICTC from administrative records from January-2009 to December-2015. To maintain confidentiality, all personal identifers, including name, address and phone numbers were delinked from the analysis dataset, by not entering them into the database, while abstrating information from the clinic register. The data was entered in Epi Info v.7, and analysis was done in Stata, v.13, College Station, Texas. Initially, descriptive analysis was done for variables on age, sex, route of transmission, followed by cross tabulation. For serodiscordant analysis, the primary client (*person reporting first*) was considered as the index case. HIV status was confirmed as per the HIV testing guidelines of the National AIDS Control Organization of India. (4)

## Results

From January 2009 to December 2015, more than 20,000 clients availed the HIV testing and counseling facility (Table 1). A total of 245 HIV positive cases were identified, of which, 148 were females (and one TG, not shown in table1). The overall HIV sero-positivity rate was 0.87%. The HIV sero-positivity rate among female clients remained uniform at around 0.5% (figure 1), while the last three-year average (2012-2014) for males was three times that of females at 1.7%. The figures 2–4 provide the age, sex, occupation and education profile of the clients.

**Table 1:** HIV Sero-positivity among clients attending ICTC at sub district hospital, Ballabgarh, 2009-2015

| **Year** | **Female Tested No. (%)** | **Male Tested No. (%)** | **Total tested** | **Number of Seropositive Female (%)** | **Number of Seropositive Male (%)** | **Total seropositivity Rate (%)** |
| --- | --- | --- | --- | --- | --- | --- |
| **2009** | 821 | 572 | 1,393 | 4 (0.5) | 7 (1.2) | 0.79% |
| **2010** | 1,256 | 833 | 2,089 | 5 (0.4) | 7 (0.8) | 0.57% |
| **2011** | 1,917 | 1,015 | 2,932 | 9 (0.5) | 22 (2.1) | 1.02% |
| **2012** | 2,767 | 1,381 | 4,148 | 15 (0.5) | 22 (1.6) | 0.89% |
| **2013** | 3,901 | 1,805 | 5,706 | 22 (0.5) | 29 (1.7) | 0.89% |
| **2014** | 4,649 | 1,909 | 6,558 | 24 (0.5) | 34 (1.8) | 0.88% |
| **2015** | 3,085 | 1,123 | 5,208 | 17 (0.5) | 27 (1.5) | 0.33% |
| **Total** | **15,311** | **7,515** | **22,826** | **96 (0.1)** | **148 (0.2)** | **0.87%** |

**Figure 1:**
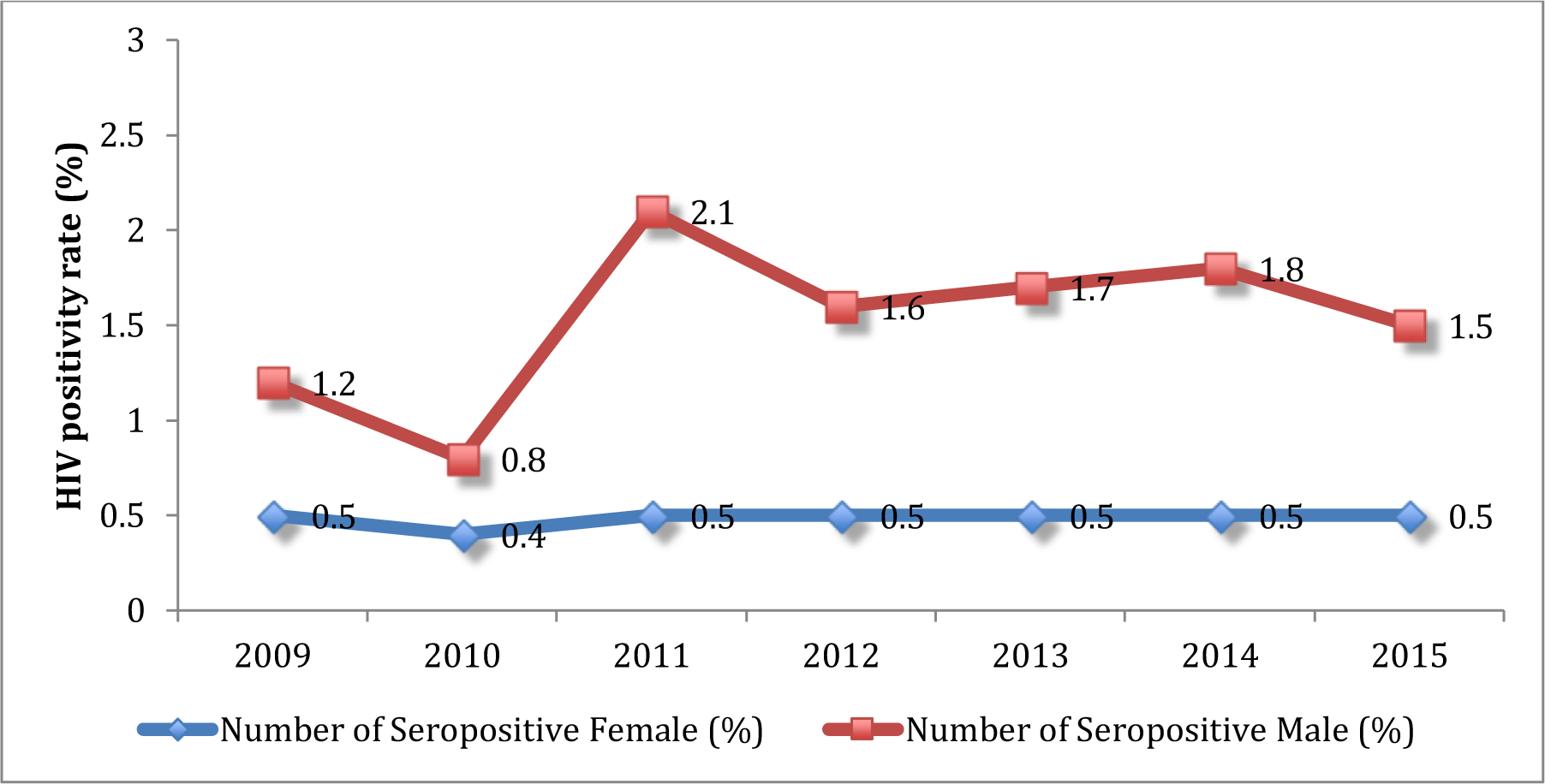
Trend of HIV sero-positivity among ICTC attendees, by sex, 2009-2015

**Figure 2:**
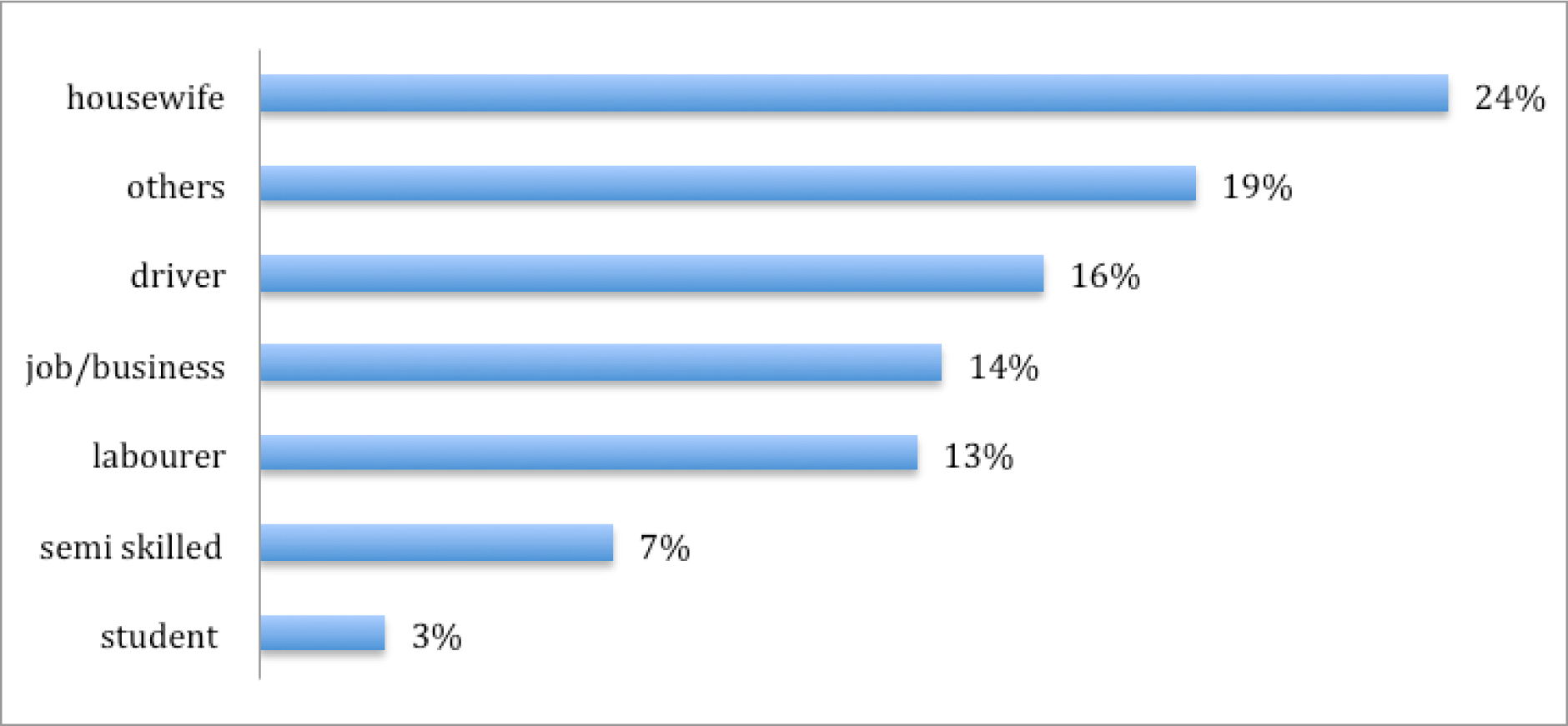
Occupation Profile of HIV positive attendees at ICTC, 2009-2015, (N=245)

**Figure 3:**
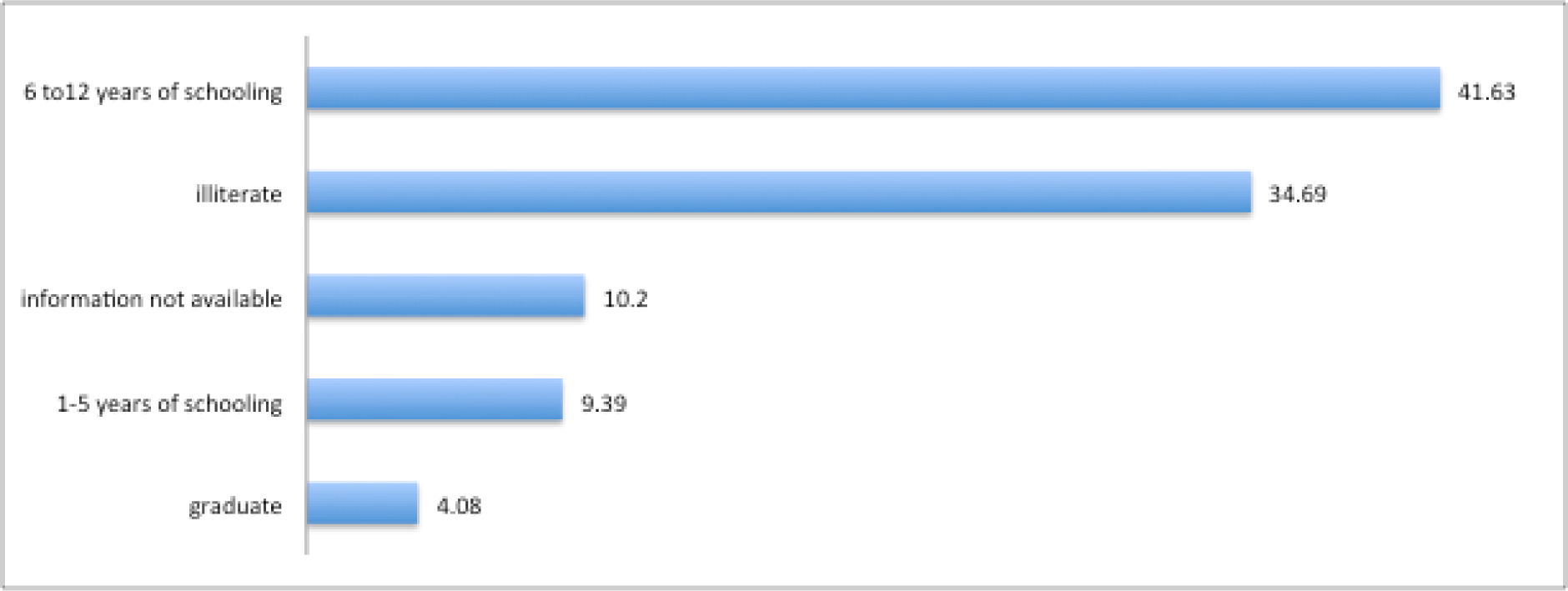
Percentage distribution of education profile of HIV positive attendees at ICTC, 2009-2015, (N=245)

**Figure 4:**
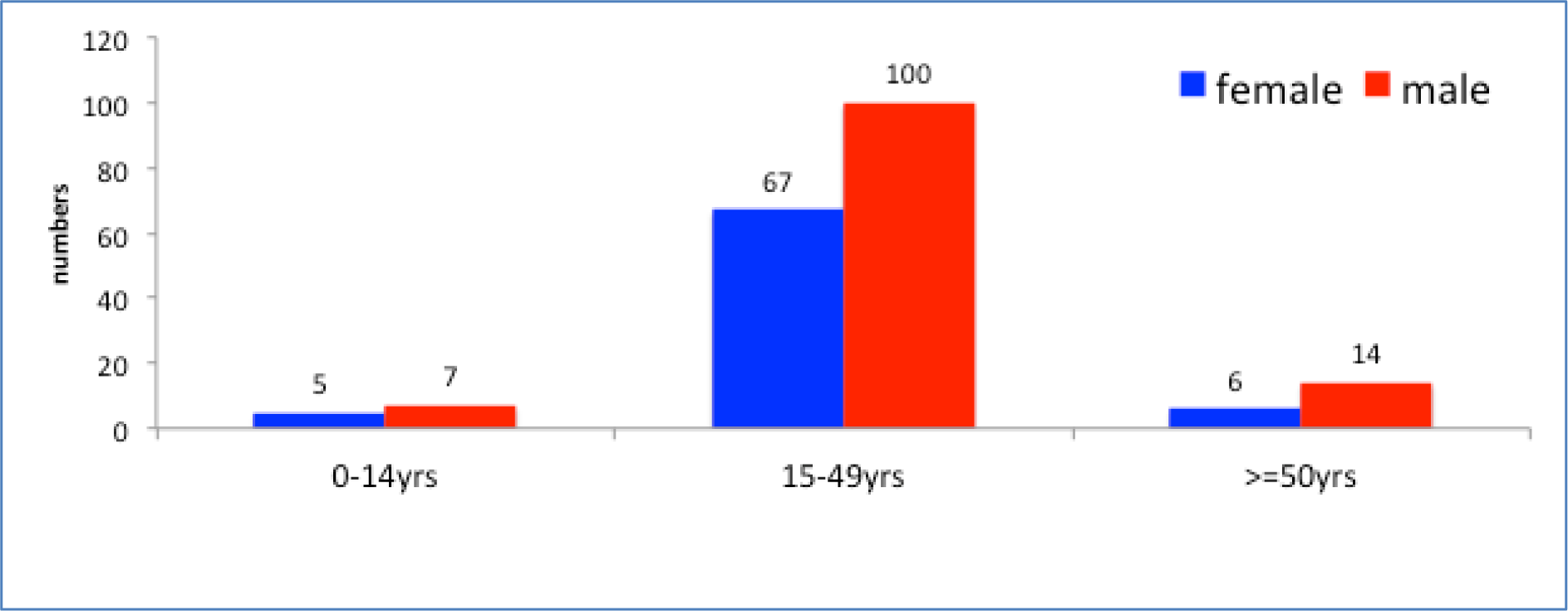
Age and Sex profile of HIV sero-positive attendees, 2009-2015

Out of 245, 39% were females. Age group of 15–49 years accounted for 82% of the total attendees, while 6% were less than 15 years of age. As shown in figure 5, the major route of transmission was heterosexual (80%). One-fifth (20%) of the clients were illiterate; 80.4% were married; and 97.7% were living with families. Most of the clients were direct-walk-in (36%), while one third were referred from Government facility. The latter comprised mainly the antenatal cases that were seen at the same health care facility. One fifth of the cases visited as part of the HIV-tuberculosis cross referral from the adjoining tuberculosis unit situated in the same health care facility (figure 6).

**Figure 5:**
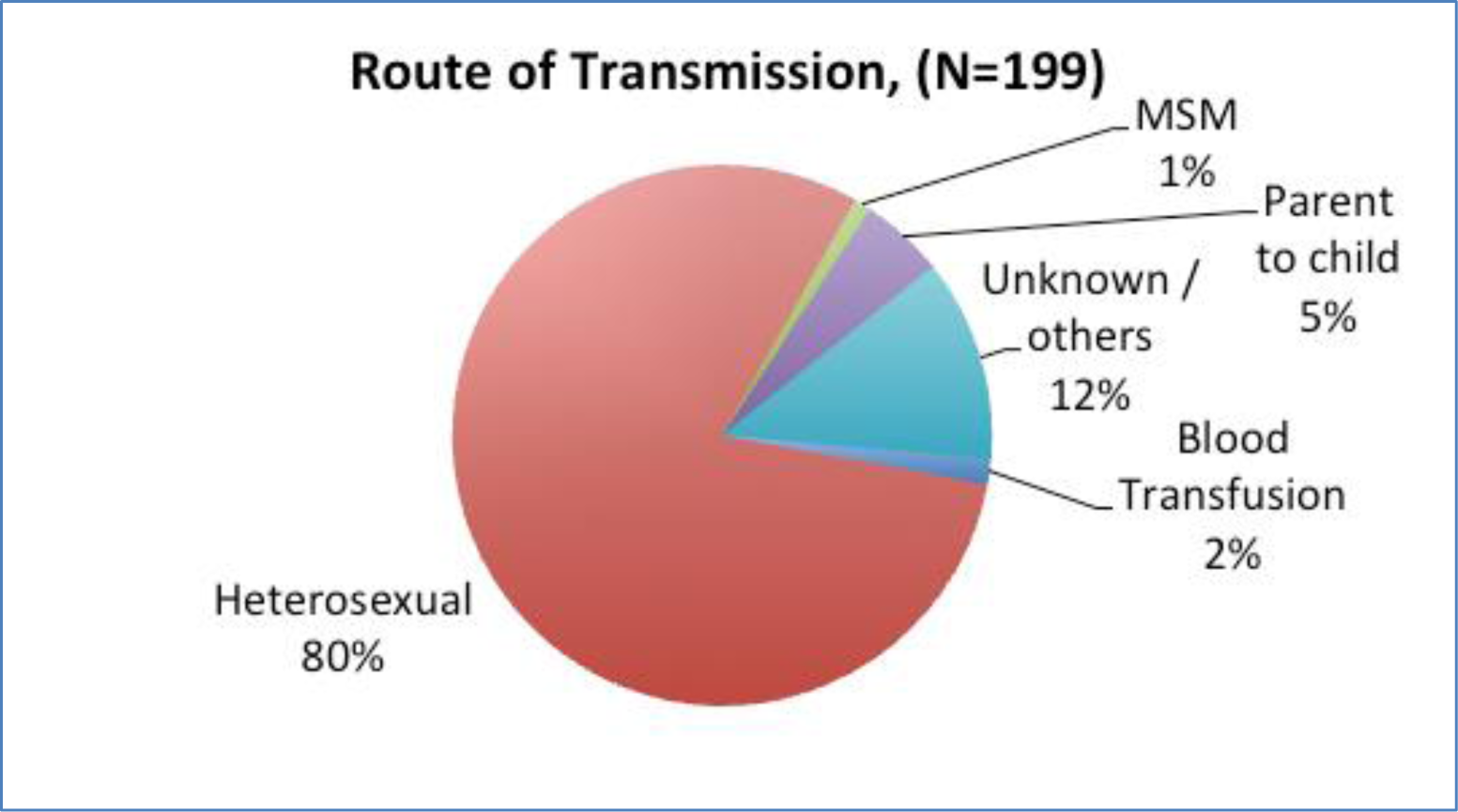
Route of Transmission, cumulative, 2009-2014

**Figure 6:**
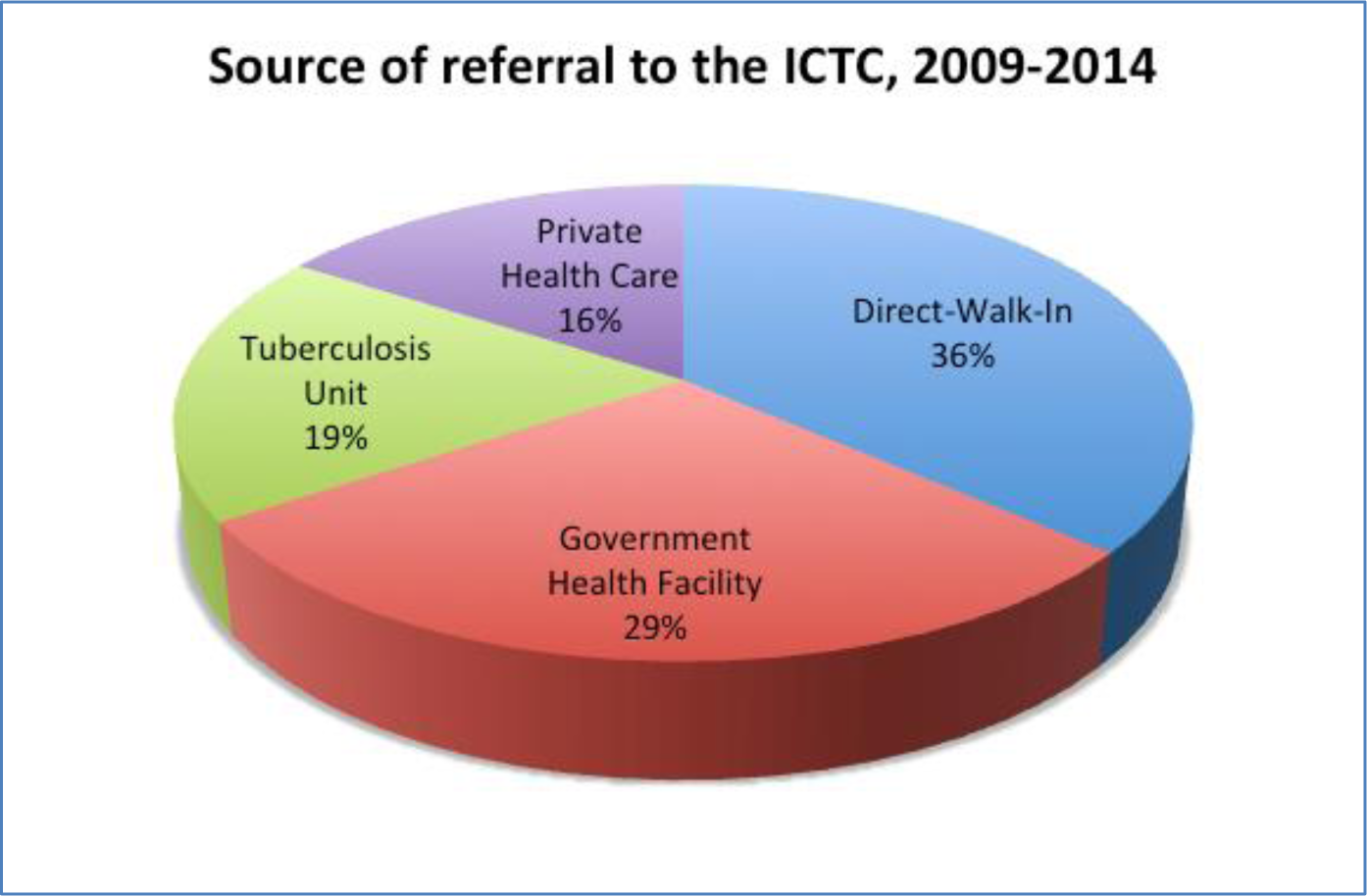
Source of referral to ICTC, cumulative 2009-2014

A total of 78 individuals, were considered index cases for the sero-discordant analysis, based on which, the analysis of 78 pairs is shown in table 2. They were 31 female, and 48 male index cases. On HIV testing of their spouses, both the spouses were positive among 45 couples, giving an overall concordance rate of 57.7%. In the remaining 33 couples, one of the spouse was negative, resulting in an overall sero-discordance rate of 42.3% in this study population. The HIV test result of their spouses further showed that when the index case was a male, the sero-discordant rate was 43.8%, and when index case was a female, it was 38.7% (table 2).

**Table 2:** HIV Sero discordance rate, cumulative, 2009-2015

| **Index Case** | **HIV Negative (%)** | **HIV positive (%)** | **Total** |
| --- | --- | --- | --- |
| **Males** | **21 (43.8%)** | 26 (54.2%) | **48** |
| **Females** | **12 (38.7%)** | 19 (61.3%) | **31** |
| **Total** | 33 (42.3%) | 45 (57.7%) |  |
|  | *Sero-dis-concordant* | *Sero-concordant* | **N=78** |

## Discussion

The Integrated Counseling and Testing Centre (ICTC) is an entry point to care and support services that provide people with an opportunity to know and understand HIV serostatus in a confidential manner. Our analysis showed that data generated at an ICTC can provide crucial information to understand the key aspects of HIV epidemiology in a particular region.

We found a total sero-positivity rate of 0.87%. The estimated HIV prevalence rate among the general population for the year 2015, in Haryana state was 0.15%.(1) As in our study, the district epidemiological profiling commissioned by NACO, Government of India, in its 2013 report, showed that the HIV prevalence among ICTC attendees (as of 2011) was low among male (2.01%) and female (1.88%) in the district of Faridabad, to which our sub district level hospital belonged.(6) Further, in line with our findings, the report also presented a stable trend of the figures.

Arora V et al reported a HIV prevalence rate of 5.3% among attendees of the ICTC from North West Delhi, which is around 50 kms from our hospital. Another recent study from a tertiary care hospital from Central Delhi reported a prevalenceof 6.3% among the attendees.(7) The reason for much higher prevalence may be because of the nature of the their facility that was a tertiary referral care, compared to our sub district level hospital.(8) The HIV counseling and tetsing services at a facility cater to those who coming either their own volition as direct walk-in, or through referral of care provides of other speciality and routine referrals from the targeted interventions by NGOs running in the area. Thus, the profile of attendees depends upon the population characteristics of the catchment area.

Similar to our studies, Sharma et al, in their study on profile of ICTC attendees in Ahmedabad Gujarat, India reported that most of the clients were in the age groupo 15–49 years.(9)

Considerable sero-discordance was reported in our study, implying prevention of transmission of HIV to the HIV negative partner is crucial. With close to one fifths of male spouses (17%) being negative, it can be said that it is not always true that males are the only ones bringing their HIV inefection to their female spouse. The latter fact becomes more important because of Haryana being a low prevalence states. More than one third of the female partners (44%) were HIV negative, which indicated that early testing of males can prevent significant number of their spouses getting HIVinfected. We found 38.7% of the females had a spouse who was HIV negative. This fact was important in the light of the evidence showed from the NACO size estimations that in the district of Faridabad, of the FSWs, the majority were home based (46.25%) FSWs.(6)

We report a HIV sero discordance rate of 42.3% and a sero concordance rate of 57.7% among couples tested for HIV in the integrated counseling and testing centre of a subdistrict hospital in Haryana, India. Mehra B et al(10) from their study in a tertiary care teaching hospital in Delhi, reported a seroconcordance rate of 45.4% and a serodiscordance rate of 54.6% among HIV-affected couples. Marfatia et al reported serodiscordanc rate of 40% in their study from Vadodara in the State of Gujarat in India.(11) These were only two studies available in literature that have reported on sero-discordance among couples from north India. The findings of our study, perhaps the third one reporting HIV sero discordance rate from north India, found the rates to be similar with these earlier two studies. However, the merit of our study is posed in two aspects. We could reaffirmed the similar sero discrodance rate as had been reported earlier, and second, ours is the first study to report HIV concordance rate from a sub district level hospital, because the other two studies were based at ICTC of tertiary care teaching hospitals.

The substantial contribution of serodiscordant partnerships to the burden of HIV/AIDS in North India highlights the fact that HIV-negative individuals in such partnerships are continually exposed to the virus and this group perhaps constitutes a major target for.

The state of Haryana, where our study was done is a low HIV prevalence area, with an estimated 1236 new HIV infections in 2015.(1) Gujarat on the other hand is showing an increasing HIV prevalence trend, as well as is contributing an estimated 10,589 new HIV infections in 2015.(1)

Several novel methods are available for prevention of HIV transmission among sero-discordant couples, such as voluntary counseling and testing, condom promotion and couple based risk reduction counseling are the foremost interventions for an effective prevention plan among discordants. (12) However, critical factors affecting the success of such prevention rely on ability of the partner to disclose their HIV status, willingness of the couple to have a regular and open communication about prevention methods including use of condom, confortness in discussing issues with having sex, and ability to refuse sex if their partner did not want to use a condomand. McGrath JW et al demonstrated feasibility of a group-based couples intervention to increase condom use in HIV serodiscordant couples in three countries (India, Thailand and Uganda), with unique HIV epidemics.(13)

There is paucity of information on HIV epidemiology at sub-district level in a low HIV prevalence State of India. Appropriate programmatic intervention for HIV Care and Treatment in a low prevalence state requires local level analysis of programme data. Heterosexual is the major mode of transmission where targeted intervention is required. Further studies are needed to explore risk factors associated with HIV transmission in discordant couples. Analysis of programme data at ICTC level is able to highlight crucial information for local HIV control.

